# Spatiotemporal Patterns of Active Deformation Reveal Downregulation of Cell-Cell Adhesion in Patient-Derived Colorectal Cancer Organoids with *BRAF* Mutations

**DOI:** 10.64898/2026.03.07.710277

**Authors:** Shogo Nagai, Ryo Suzuki, Go Yamakawa, Akihisa Fukuda, Hiroshi Seno, Motomu Tanaka

## Abstract

Colorectal cancer (CRC) is the second most common cause of cancer-related mortality. At the molecular level, CRC is associated with genetic mutations and epigenetic modifications that dysregulate various signaling networks. From the biophysical viewpoint, invasive and metastatic cell migration need to be empowered by mechanical forces. In this study, we analyze the dynamic deformation of patient-derived CRC organoids in Fourier space and demonstrate how organoids with protooncogene *BRAF* mutation exhibit deformation phenotypes at an early stage. The organoids with *BRAF*^mut^ have significantly lower elasticity and higher viscosity than those with *BRAF*^WT^, which mathematically indicated as the weakening of cell–cell adhesion. Immunohistochemical images, qRT-PCR, and TCGA data analysis confirm the downregulation of E-cadherin (*CDH1*) in *BRAF*^mut^ organoids as well as in *BRAF*^mut^ CRC, suggesting that the decrease in cell–cell adhesion in *BRAF*^mut^ CRC facilitates invasive and metastatic migration. Notably, the recovery of *CDH1* expression by pharmacological inhibition of DNA methylation can quantitatively be detected as the change in mechanical properties, suggesting that the complementary combination of dynamic phenotyping, mathematical modelling, and molecular-level analyses has a potential to unravel the mechanistic causality of the critical gene mutation and CRC’s prognosis and the response to therapeutic interventions.

## 1. Introduction

Colorectal cancer (CRC) is the third most common cancer worldwide and the second most common cause of cancer-related death. It is often diagnosed at an inoperable stage after local invasion and distant metastases [1]. Moreover, CRC often develops resistance to conventional chemotherapy. CRC is known to be associated with mutations of several major oncogenes and tumor suppressor genes, such as *APC*, *KRAS*, *BRAF* and *TP53*. In addition to such gene mutations, recent studies have demonstrated the involvement of epigenetic modifications in CRC, including DNA methylation, histone modification and chromatin remodeling [2]. Genetic mutations and epigenetic modifications cause dysregulation of conserved signaling networks, exerting context-dependent effects on critical phenotypes. Among these, one of the critical mutations is that of proto-oncogene *BRAF*, which is present in 8%–15% of CRC cases [3]. *BRAF* is a protooncogene involved in the EGFR pathway, located downstream of *KRAS* and upstream of *MEK*. A recent study showed that SRC kinase promotes the resistance to BRAF/EGFR targeted therapy [4]. *BRAF* driver mutation, represented by *BRAF^V600E^*, is one of the characteristics of consensus molecular subtype 1 (CMS1) [5] that shows a wide-spread epigenetic DNA methylation. Because *BRAF* mutation in CRC frequently results in poor prognostic outcomes, it is clinically important to establish the appropriate management of CRC with *BRAF* mutation [6].

Over the last two decades, three-dimensional (3D) cancerous tissues cultured *in vitro*, called cancer organoids, have been used as an alternative to the conventional two-dimensional (2D) culture of cell lines. Organoids are generally defined as microscopic, self-organizing, three-dimensional structures, which can be grown from stem cells and self-organize into 3D structures *in vitro* [7]. Patient-derived organoids established from various cancerous tissues have been attracting increasing attention because they are expected to provide more information about the efficacy of different treatments due to their phenotypic and genotypic similarity to the tumors of origin [8]. Codrich et al. performed genomic, transcriptomic and phosphoproteomic analyses of patient-derived CRC organoids and demonstrated that they recapitulate the original tumors at the molecular level, suggesting that they can be used as a 3D model mimicking the evolution of the patient’s tumor *in vivo* [8a]. Multi-omics analysis and microscopy-based phenotypic observation of fixed organoids have contributed to unravel how genetic mutations or epigenetic cues modulate the signaling pathways and the characteristic patterns of protein expression in various cancer types [9].

Recently, an increasing number of studies have shown the complex interplay between mechanical forces and biochemical signals in regulating dynamic changes in tissue geometry during morphogenesis associated with development and diseases [10]. At the single-cell level, forces are generated either by contraction of the actomyosin complex or by the polymerization of cytoskeletal proteins, which modulates the mechanical properties of cells. At the multicellular scale, these forces are transmitted to the extracellular matrix and neighboring cells via adhesion molecules, such as integrin and cadherin. Several mathematical models have been developed to reproduce the formation of 3D intestinal organoids observed experimentally [11]. For example, Buske et al. developed a model of crypt formation and intestinal organoid formation, in which a basement membrane was modelled as a triangular lattice network of “stiff” polymers and the cells as elastic spheres adhering to the underlying basal membrane [12]. Cells in cancer organoids not only exhibit large-scale rearrangements and proliferation over several days but also undergo a variety of dynamic deformations on the timescales of minutes to hours. In the context of CRC, the mechanistic link between genetic and epigenetic alterations and the spatiotemporal regulation of active deformation of multicellular systems – and its role in shaping complex tissue architectures – is poorly understood. Understanding this relationship is crucial, because these active forces influence tissue morphogenesis, leading to invasive and metastatic characteristics of tumors.

In this study, we investigate how patient-derived CRC organoid with critical oncogene mutation modulates the fast deformation forces. The impact of *BRAF* driver mutation, which frequently results in poorer prognoses in CRC patients, was a particular focus of this work. Dissociated CRC cells were seeded into a gel matrix, and the dynamic deformation was monitored by time-lapse imaging every 10 min from the single-cell stage up to *t* ≈ 80 h. Fourier mode analysis of active deformation indicated that the shape relaxation after the cell division of *BRAF*^mut^ was markedly slower than that of *BRAF*^WT^. The experimental data were fitted with the classical Voigt model, highlighting that organoids with *BRAF* mutation significantly modulated the effective elasticity and viscosity of organoids. With the aid of mathematical modelling of the gradient flow of free energy, the experimentally observed change in viscoelastic shape recovery could be explained by the modulation of intercellular tension. Intriguingly, direct comparison of the immunohistochemical images of *BRAF*^mut^ and *BRAF*^WT^ organoids indicated that they differed markedly in the expression of E-cadherin and F-actin, proteins that contribute to cell–cell adhesion and force generation. qRT-PCR analyses of *BRAF*^mut^ and *BRAF*^WT^ further demonstrated the significant downregulation of *CDH1* in *BRAF*^mut^.

Moreover, the differential expression of E-cadherin between *BRAF*^mut^ and *BRAF*^WT^ in *in vivo* tissues was confirmed by immunohistological staining of the original CRC tissues and large-scale PanCancer Atlas cohort data in The Cancer Genome Atlas (TCGA), which molecularly characterized over 20,000 primary cancers and matched normal samples from 33 cancer types [13]. The PanCancer Atlas cohort analysis suggests that epigenetic DNA methylation contributes to the downregulation of *CDH1* in *BRAF*^mu^t CRC. In fact, the pharmacological inhibition of epigenetic DNA methylation restored *CDH1* expression in a concentration-dependent manner and reverted the dynamic phenotype of *BRAF*^mut^ organoids to that of *BRAF*^WT^. These data indicate that epigenetic regulation in *BRAF*^mut^ organoids modulates cadherin-mediated intercellular tension. Furthermore, cohort and pathway analyses revealed downregulation of the canonical WNT pathway and upregulation of planar cell polarity (PCP) signaling that might promote invasive and metastatic cell migration characterizing *BRAF*^mut^ CRC. Our study demonstrates that dynamic phenotyping of CRC organoids offers a quantitative approach to connect genetic/epigenetic alterations, therapeutic interventions, and prognostic phenotypes.

## 2. Results

### 2.1. Spatiotemporal dynamics of patient-derived CRC organoids

As schematically presented in **Figure 1a**, the dissociated cells in patient-derived CRC organoids were seeded in Matrigel, and the spatiotemporal dynamics of active deformation was recorded by label-free, brightfield time-lapse imaging from the single-cell stage up to *t* ≈ 80 h. This time window was chosen in order to focus on the active deformation of early-phase CRC organoids before cyst formation (Figure S1, Supporting Information). The characteristics of the patients are summarized in **Table 1**. **Figure 1b** shows the brightfield images of *BRAF*^WT^ (C43T) and *BRAF*^mut^ (C7T) captured at *t* = 0, 20, 40, 60 and 80 h. The original time-lapse movies are presented in Movie S1 and S2, Supporting Information. The average number of cell nuclei counted at *t* ≈ 80 h was 11.7 ± 3.1 for *BRAF*^WT^ (*N* = 35) and 12.0 ± 2.8 for *BRAF*^mut^ (*N* = 26), suggesting that most cells underwent three to four divisions during this period (Figure S2, Supporting Information). The images were acquired every 10 min, and the outer rim (contour) in the 2D projection was extracted for each frame using machine learning (**Figure 1c**). The complex deformation was analyzed by Fourier transformation of the radial distance between the contour and the center of mass *r*(*θ*, *t*) [14]:

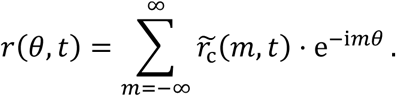

**Figure 1.**
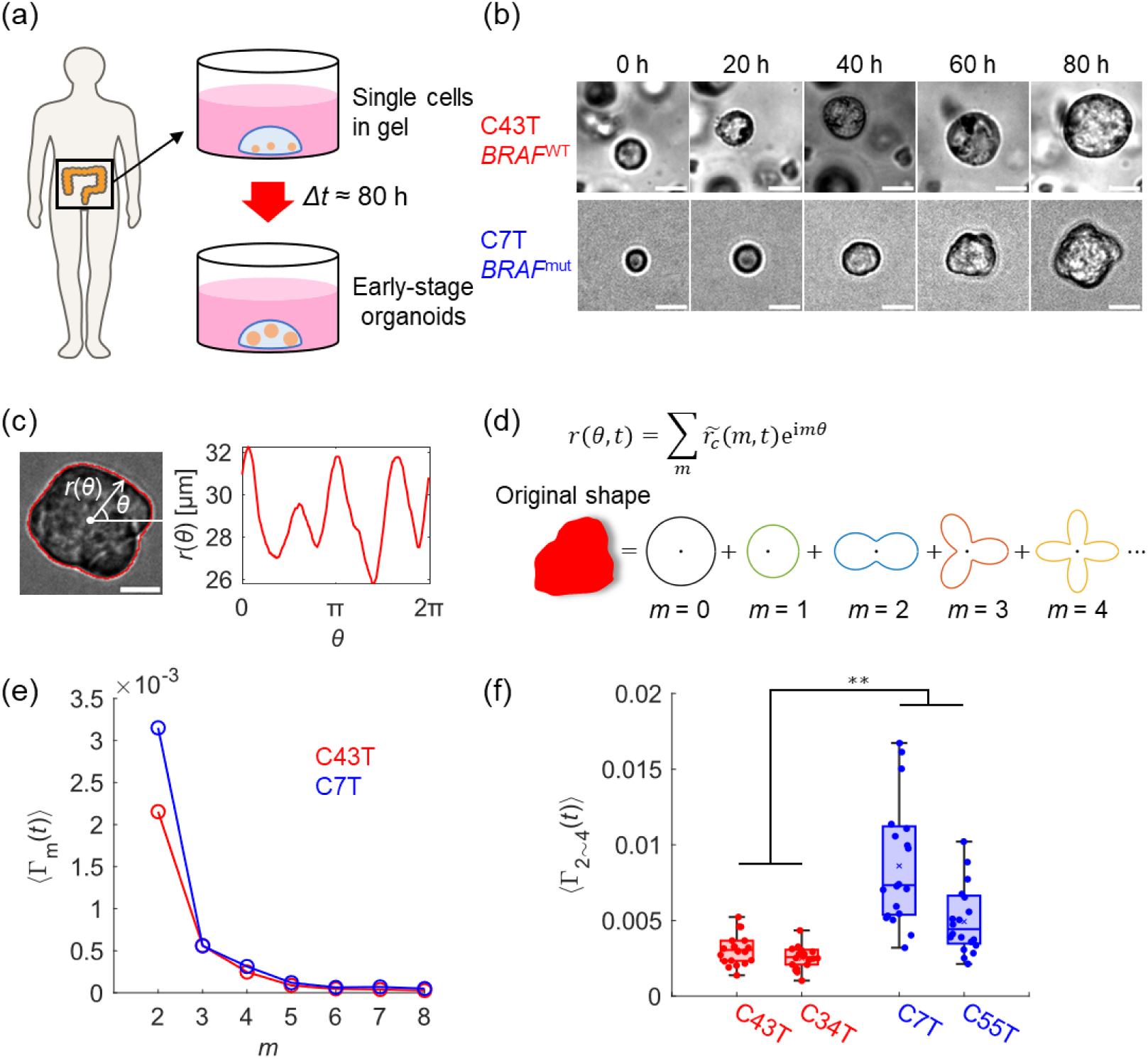
Spatiotemporal dynamics of patient-derived colorectal cancer organoids. (a) Dynamic deformation of patient-derived colorectal cancer organoids embedded in Matrigel was monitored from the single-cell stage up to *t* ≈ 80 h, when no hollow cyst formation could be detected (Figure S1). (b) Snapshot images of organoids derived from patients with wild-type *BRAF* (*BRAF*^WT^) and those with *BRAF* mutation (*BRAF*^mut^). (c) Radial distance from centroid *r*(*θ*, *t*) was extracted from each time-lapse image. (d) Complex deformation was evaluated by Fourier mode analysis. (e) Power spectra of *BRAF*^WT^ (C43T, red) and *BRAF*^mut^ (C7T, blue) normalized by average size 〈Γ_*m*_(*t*)〉. No marked difference could be detected for *m* ≥ 5. (f) *BRAF*^mut^ (C7T, *N* = 19; C55T, *N* = 19, blue) averaged over *t* = 0 ∼ 80 h exhibited a significant difference in total deformability 〈Γ_2∼4_(*t*)〉 than *BRAF*^WT^ (C43T, *N* = 19; C34T, *N* = 18, red). **: *p* < 0.01. Scale bars: 20 µm.

**Table 1.** Summary of patient characteristics.

|  | Organoid# | Age | Sex | Location | Histology | Stage |
| --- | --- | --- | --- | --- | --- | --- |
| *BRAF*<sup>WT</sup> | C34T | 80 | F | Transverse<br>colon | Moderately<br>differentiated | II |
|  | C43T | 77 | M | Ascending<br>colon | Well<br>differentiated | I |
| *BRAF*<sup>mut</sup><br>*V600E mutation | C7T | 64 | M | Transverse<br>colon | Mucinous | I |
|  | C55T | 91 | F | Ascending<br>colon | Moderately<br>differentiated | III |

As schematically shown in **Figure 1d**, *m* ≥ 2 represents anisotropic deformation. **Figure 1e** shows the power spectra of *BRAF*^WT^ (C43T, red) and *BRAF*^mut^ (C7T, blue), normalized by the average size:

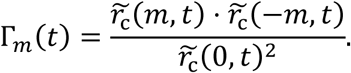

The observed deformation originates from active processes such as membrane deformation and cytoskeleton remodeling, which accompanies the dissipation of energy [14c, 15]. As no marked difference could be detected for *m* ≥ 5 in the power spectra of both *BRAF*^WT^ (C43T, red) and *BRAF*^mut^ (C7T, blue), the time-averaged total deformability summed over *m* = 2 ∼ 4:

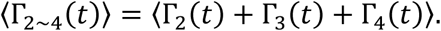

**Figure 1f** shows the total deformability of 2 × *BRAF*^WT^ (C43T and C34T, red) and 2 × *BRAF*^mut^ (C7T and C55T, blue) averaged over *t* = 0 ∼ 80 h. The statistical comparison of 〈Γ_2∼4_(*t*)〉 values indicated that the deformation of both *BRAF*^mut^ organoids averaged over 80 h was significantly larger than that of both *BRAF*^WT^ organoids (*p* < 0.01), irrespective of the cancer stage and location.

### 2.2. Viscoelastic recovery after cell division confers mechanical properties of organoids

Note that the power spectra and total deformation level presented in **Figure 1e** and **1f** were averaged over 80 h, which allows three to four cell divisions. It is well established that the event of cell division is accompanied by significant deformation. Particularly for early-phase organoids consisting of a small number of cells, cell division has a prominent effect on the dynamics. After such division, the cells undergo viscoelastic relaxation towards a sphere, or a circle in 2D projection, by minimizing the free energy corresponding to tension. Thus, to gain further insights into the viscoelastic recovery, we monitored Γ_2∼4_(*t*) every 10 min for *BRAF*^WT^ (C43T, red) and *BRAF*^mut^ (C7T, blue) organoids. **Figure 2a** shows the time-course of Γ_2∼4_ at *t* = 0 ∼ 80 h. The spike-like peaks that appear every Δ*t* = 20 ∼ 30 h correspond to individual cell divisions, while the interval between them coincides with the cell cycle. Note that the first two peaks, indicated by *t*_1_and *t*_2_ for each organoid, are more prominent than the following peaks because the deformation was normalized by the average size. **Figure 2b** presents magnified views of the first Γ_2∼4_(*t*) peaks of *BRAF*^WT^ (C43T, red) and *BRAF*^mut^ (C7T, blue). As the brightfield images indicate, a round single cell divided into two cells at *t* = *t*_1_, and the shape of the cell dimer gradually became round over time. The relaxation of deformation Γ_2∼4_(*t*) at *t* ≥ *t*_1_ can be fitted with an exponential function, Γ_2∼4_ = *A* exp(− *t*⁄*τ*) + *c* (solid line), yielding the characteristic relaxation time for *BRAF*^WT^ (C43T, red) and *BRAF*^mut^ (C7T, blue), *τ*_C43T_ = 0.2 h and *τ*_C7T_ = 0.8 h, respectively. **Figure 2c** presents statistical comparisons of the *τ* values. Both *BRAF*^mut^ organoids (C7T and C55T) showed significantly slower relaxation than both *BRAF*^WT^ organoids (C43T and C34T) (*p* < 0.01). To obtain deeper insights into the observed viscoelastic relaxation, we calculated the effective elasticity *k* and viscosity *η* of CRC organoids using the classical Kelvin–Voigt model (**Figure 2d**), the relationship between active stress *σ*(*t*) and strain *ε*(*t*) is given by:

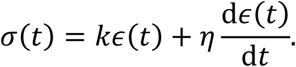

**Figure 2.**
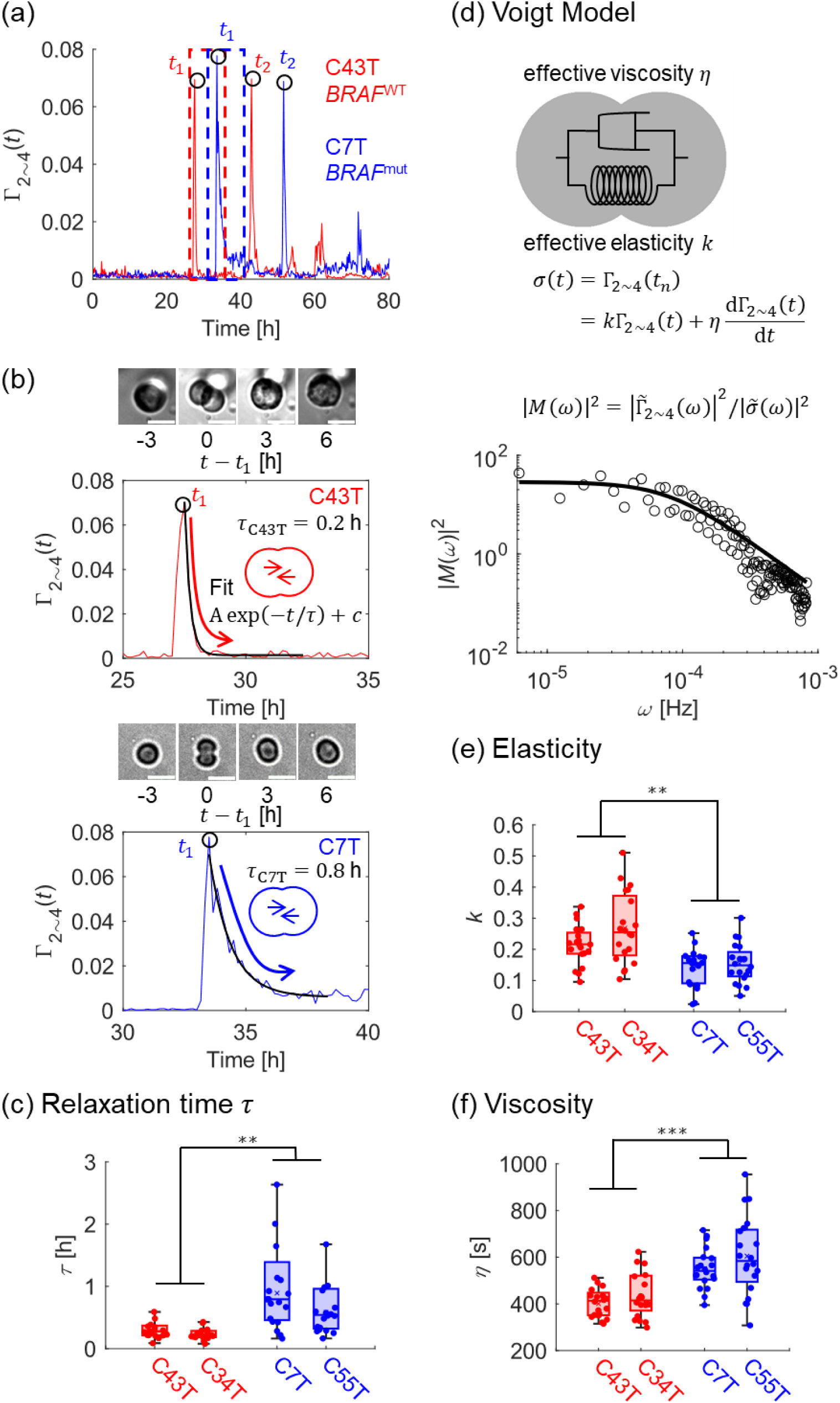
Viscoelasticity of organoids calculated from shape recovery after cell division. (a) Time-course of total deformability Γ_2∼4_ plotted as a function of *t*. Prominent peak *t*_*n*_ coincides with the timepoint of the *n*th cell division. (b) Expanded plot of Γ_2∼4_ vs. *t* for *BRAF*^WT^ (C43T, red) and *BRAF*^mut^ (C7T, blue). The dynamic shape recovery after *t*_1_ can be fitted well with an exponential function, Γ_2∼4_(*t*) = *A* exp(− *t*⁄*τ*) + *c* (black lines; *τ*_*C*43*T*_ = 0.2 h; *τ*_*C*7*T*_ = 0.8 h). (c) Statistical comparison of characteristic relaxation time *τ* of *BRAF*^WT^ (red; C43T, *N* = 18; C34T, *N* = 19) and *BRAF*^mut^ (blue; C7T, *N* = 18; C55T, *N* = 18). The recovery of *BRAF*^mut^ was significantly slower than that of *BRAF*^WT^. (d) Classical Voigt model used to calculate the effective elasticity *k* and viscosity *η*. |*M*(*ω*)|^2^is the ratio of the power spectrum of strain and that of active stress. Calculated elasticity *k* (e) and viscosity *η* (f) for *BRAF*^WT^ (red; C43T, *N* ≥ 19; C34T, *N* ≥ 19) and *BRAF*^mut^ (blue; C7T, *N* ≥ 19; C55T, *N* = 20). **: *p* < 0.01, ***: *p* < 0.001. Scale bars: 20 µm.

Here, the force to divide the cells is the active stress *σ*(*t*), while the total deformation Γ_2∼4_(*t*) is the strain *ε*(*t*). In the frequency domain, this relationship can be written by Fourier transformation:

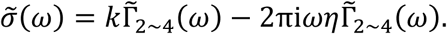

The ratio between the power spectrum of strain 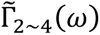 and active stress 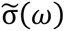 can be used to calculate the elasticity *k* and viscosity *η*:

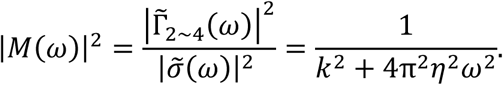

**Figure 2d** presents a double logarithmic plot of |*M*(*ω*)|^2^as a function of *ω*, and the solid line represents the fit. The calculated elasticity *k* and viscosity *η* values of *BRAF*^WT^ organoids (C43T and C34T, red) and *BRAF*^mut^ organoids (C7T and C55T, blue) are presented in **Figure 2e** and **2f**, respectively. The *BRAF*^mut^ organoids possess significantly lower elasticity and higher viscosity than the *BRAF*^WT^ ones, implying that *BRAF* mutation significantly modulates the mechanical properties of organoids well before the formation of hollow cysts.

### 2.3. Mathematical model demonstrates that *BRAF^mut^* organoid possesses altered intercellular tension

The modulation of viscoelasticity of early-phase organoids suggested by the spatiotemporal analysis implies that the transmission of force between neighboring cells is altered in organoids with *BRAF* mutation. To verify this, we simulated the viscoelastic relaxation after the first cell division (*t* ≥ *t*_1_) in 2D space using the mathematical model previously used for the cellular arrangement in 3D [16]. As schematically depicted in **Figure 3a**, two adjacent cells and their surroundings were expressed as regions *C*_1_, *C*_2_and *C*_0_, respectively. The line tension acting at the boundary between the neighbouring regions *C*_*i*_ and *C*_*j*_ was defined as *T*_*ij*_, whereas the length of boundary was defined as *L*_*ij*_. Cellular deformation and rearrangement were considered as the *L*^2^-gradient flow of the total free energy in 2D space:

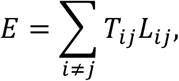

assuming that the area of each region is preserved (see Method S1, Supporting Information for more details). Here, the tension between cells varied to *T*_12_ = 1 and 12, while the tension between a cell and its surroundings was kept constant, *T*_*i*0_ = 20 (*i* = 1, 2). Note that the cell–cell adhesion becomes stronger when the intercellular tension is lower. The snapshot images of the simulation showed a clear difference in the kinetics of shape recovery at the different levels of tension between cells (**Figure 3b**). The simulation movies are presented Movie S3 – S5, Supporting Information. When the intercellular tension (*T*_12_ = 1) is lower than the tension between cells and their surroundings (*T*_*i*0_ = 20), two cells quickly recover the circular shape (in 2D) to minimise the tension free energy *E* (top). In stark contrast, the recovery becomes slower when the intercellular tension *T*_12_ becomes closer to *T*_*i*0_, such as *T*_12_ = 12 (bottom). In fact, at *T*_12_ = 19, the cells hardly recover the circular shape at all (Figure S3, Supporting Information). **Figure 3c** presents the time-course of calculated deformation Γ_sim_ corresponding to *T*_12_ = 1 (red) and *T*_12_ = 12 (blue), which reproduced the total deformation Γ_2∼4_ obtained by the experiments (**Figure 3d**). Thus, our mathematical modelling provided numerical evidence that *BRAF^mut^* organoid possesses altered intercellular tension in CRC organoids, resulting in the differential relaxation kinetics.

**Figure 3.**
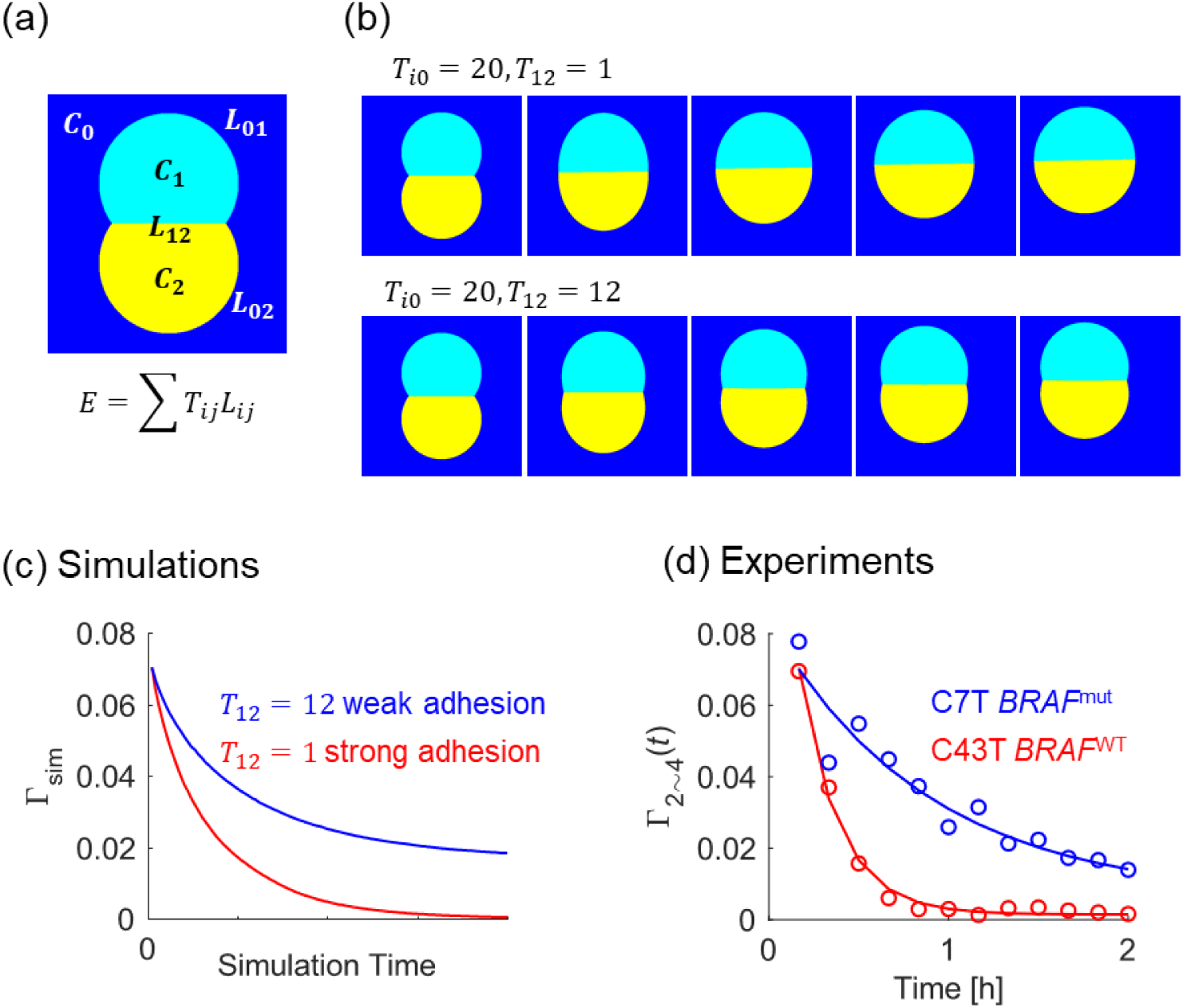
Mathematical model demonstrates that *BRAF^mut^* organoid possesses altered intercellular tension. (a) Schematics of the mathematical model. Shape recovery is simulated by gradient flow of tension energy *E* represented by line tension *T*_*ij*_ and boundary length *L*_*ij*_ between regions *C*_*i*_ and *C*_*j*_, *E* = ∑ *T*_*ij*_*L*_*ij*_. (b) Snapshot images of simulations corresponding to two different intercellular tensions *T*_12_. Calculated deformation Γ_sim_ (c) reproduces experimental data Γ_2∼4_ (d), suggesting that the modulation of intercellular tension caused by *BRAF* mutation results in the differential relaxation kinetics of CRC organoids.

### 2.4. *BRAF^mut^* organoid shows weaker cell–cell adhesion and contractile forces

In CRC organoids, the mechanical forces generated by the contraction of actomyosin are transmitted to the extracellular matrix and to neighboring cells via adhesion contacts [10c]. The mechanical interaction with the gel matrix was found to be negligible in this study because Young’s modulus was ∼100 Pa, which is well below the level for activating the molecular clutch machinery via focal adhesion [17]. To unravel how cell–cell adhesion is altered in CRC organoids with *BRAF* mutation at the protein level, we investigated the expression patterns of E-cadherin by immunohistochemistry (**Figure 4a**). *In vivo*, it is known that a compromised epithelial barrier in the colon leads to inflammation and tissue fibrosis in the context of CRC [18]. As E-cadherin molecules in cell–cell junctions are connected to F-actin via α- and β-catenin crosslinkers, we also labelled F-actin in *BRAF*^WT^ (C43T) and *BRAF*^mut^ (C7T) organoids (**Figure 4b**). The networks of F-actin myosin motors are the generators of contractile forces, and the cortical tension and its gradient determine the mechanical properties of tissues [19]. Previous reports have shown that the dysregulation of E-cadherin-mediated cell adhesion and F-actin networks resulted in the loss of contact inhibition in CRC [20]. All of the images were taken at *t* = 72 ∼ 80 h because the signal intensities from the actin concentrated near the future cyst become dominant after this timepoint (Figure S4, Supporting Information). **Figure 4c** and **4d** show the statistical comparisons of integrated E-cadherin and F-actin signals from the plane of the equator, respectively. The data extracted from *N* ≥ 10 samples indicated that the expression level of E-cadherin is significantly lower in *BRAF*^mut^ than in *BRAF*^WT^ (**Figure 4c**, *p* < 0.05). This is in line with the findings of a previous study on colon adenocarcinoma cells with ectopically expressed or silenced mutations of *BRAF*, showing significant impairment of the distributions of E-cadherin, which were characterized by noncontinuous, fragmented adherens junctions [21]. The lower E-cadherin signals in *BRAF*^mut^ agree well with the slower shape relaxation reproduced by the numerical simulations at weaker adhesion and hence higher intercellular tension, *T*_12_ = 12 (**Figure 3**). The significantly lower F-actin signals in *BRAF*^mut^ (*p* < 0.001, **Figure 4d**) appear to correlate with the impaired expression of E-cadherin, resulting in the lower elasticity *k* and higher viscosity *η* (**Figure 2**).

**Figure 4.**
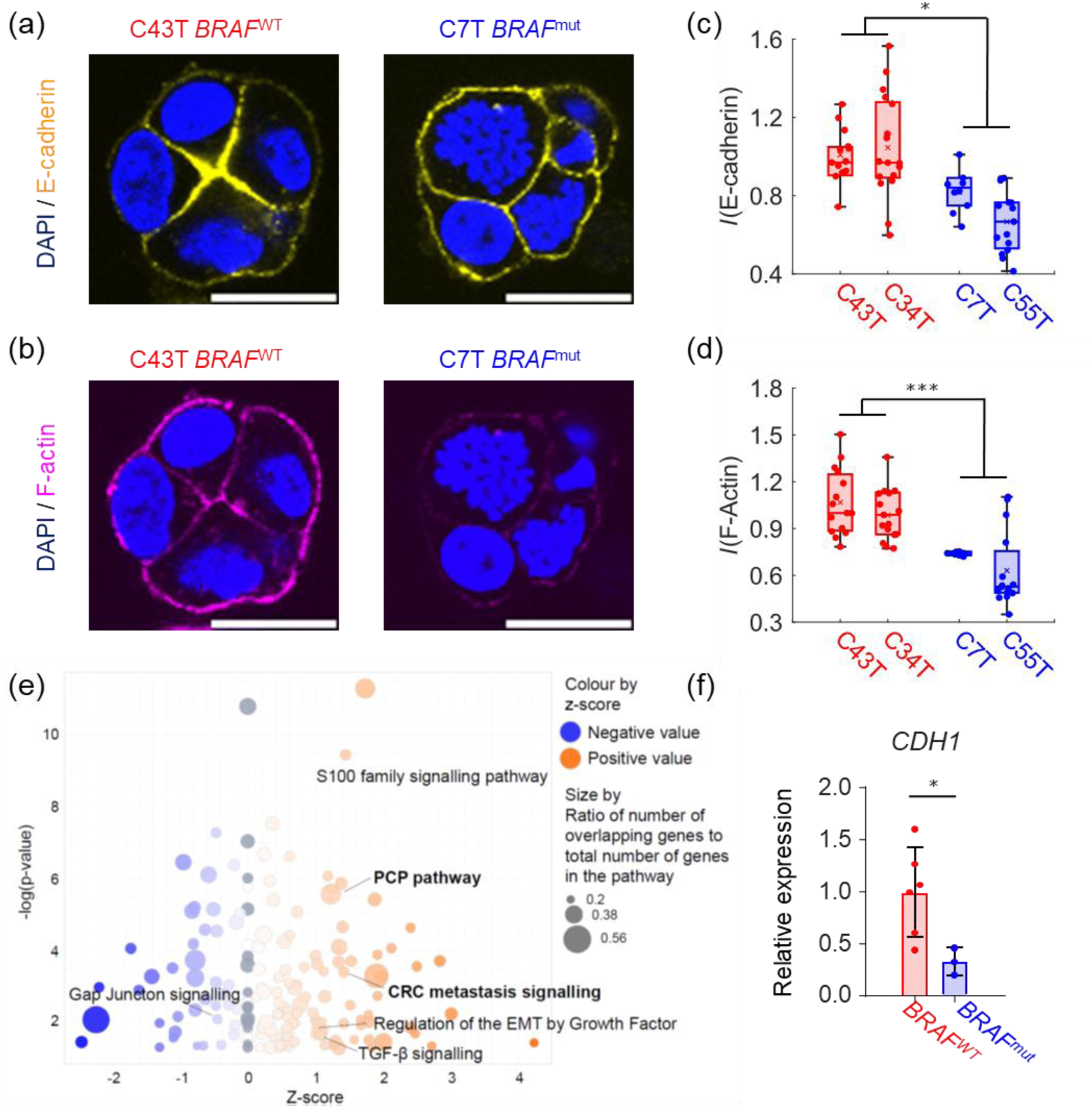
Weakening of cell–cell adhesion in *BRAF^mut^* organoids. Immunohistochemical staining of E-cadherin (yellow, a) and F-actin (magenta, b) for *BRAF*^WT^ (C43T) and *BRAF*^mut^ (C7T). Each image is a slice corresponding to the plane of the equator. Integrated signals from *BRAF*^WT^ (C43T,C34T, red) and *BRAF*^mut^ (C7T,C55T, blue) indicate that the fluorescence signal intensities of E-cadherin (c) and F-actin (d) are significantly different (*: *p* < 0.05, ***: *p* < 0.001). (e) Ingenuity Pathway Analysis. Dot plots showing the up- or downregulated pathways in *BRAF*^mut^ compared with the levels in *BRAF*^WT^. Each dot represents a significantly enriched pathway, and its size corresponds to the ratio of the number of overlapping genes to the total number of genes in the pathway. Only the pathways with FDR < 0.01 are plotted. (f) Relative expression levels of the *CDH1* gene in *BRAF*^WT^ (C43T, red) and *BRAF*^mut^ (C7T) exhibit a significant difference (*: *p* < 0.05). Scale bars: 20 µm.

We performed RNA-seq on the *BRAF*^wt^ (C34T and C43T) and *BRAF*^mut^ (C7T and C55T) organoids to identify differentially expressed genes (DEGs) with log fold change > |1|. As shown in **Figure 4e**, Ingenuity Pathway Analysis of the DEGs suggested that the migration-related PCP pathway is upregulated in *BRAF*^mut^ compared with the level in *BRAF*^WT^, suggesting the potential involvement of epithelial–mesenchymal transition (EMT) and metastasis [22]. Our immunohistochemical data (**Figure 4c**) are in good agreement with the previous studies suggesting the downregulation of E-cadherin by the modulation of pathways such as the TGF-β and WNT/β-catenin signaling pathways [23]. In fact, qRT-PCR data further confirmed that the expression of *CDH1* in *BRAF*^mut^ organoids was significantly lower than that in *BRAF*^WT^ organoids at the mRNA level (*p* < 0.05) (**Figure 4f**).

### 2.5. DNA methylation is a factor causing impaired E-cadherin expression in *BRAF*^mut^ CRC in vivo

To determine whether the differential expression levels of E-cadherin observed in organoids reflect those in CRC tissues *in vivo*, we immunohistochemically stained E-cadherin in the original tissues. **Figure 5a** shows immunohistochemical images of tissue slides from *BRAF*^WT^ and *BRAF*^mut^ CRC from which organoids were prepared. The level of E-cadherin expression in *BRAF*^WT^ tissues was higher than that of *BRAF*^mut^ tissues (**Figure 5a**, insets). The expression levels of the *CDH1* gene in *BRAF*^WT^ and *BRAF*^mut^ CRCs were statistically compared by analyzing TCGA cohort data (*N* = 311), demonstrating that the *CDH1* gene was significantly downregulated in CRC tissues with *BRAF* driver mutation (*p* < 0.001) (**Figure 5b**).

**Figure 5.**
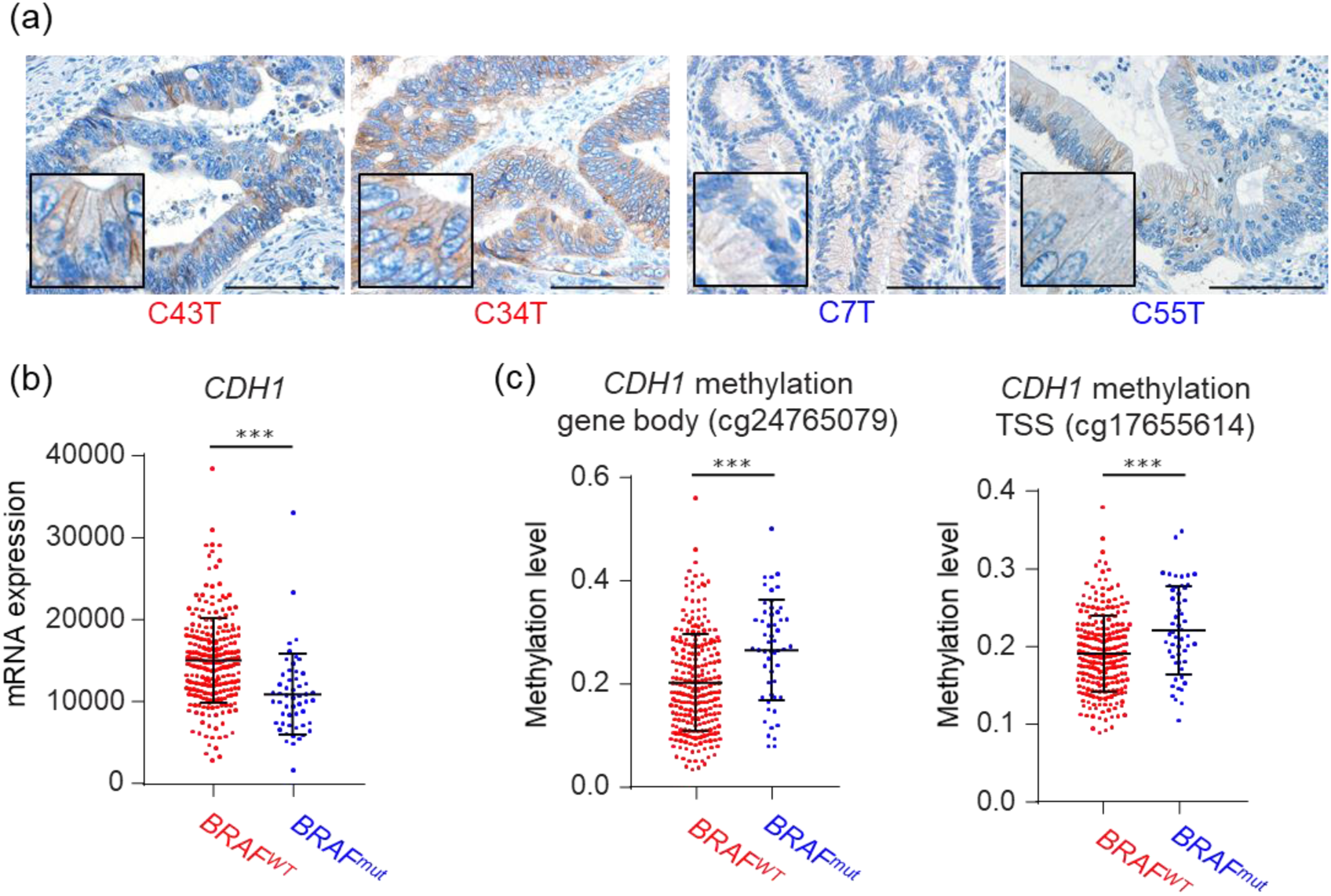
DNA methylation as a factor causing impaired E-cadherin expression *in vivo*. (a) Immunohistochemical images of the original CRC tissues of *BRAF*^WT^ (C43T, C34T) and *BRAF*^mut^ (C7T, C55T) from which organoids were established. Higher-magnification images (bottom left) suggest that membrane expression of E-cadherin is less pronounced in *BRAF*^mut^ tissue. Scale bars: 100 µm. (b) *CDH1* expression profiles obtained from TCGA datasets (*N* = 311) indicate that significant downregulation of *CDH1* in CRC tissues with *BRAF* mutation at the transcript level. (c) The PanCancer Atlas cohort data in TCGA shows higher levels of DNA methylation at two CpG sites of *BRAF*^mut^ tissues compared with those in *BRAF*^WT^, suggesting epigenetic DNA methylation is one of the key factors causing the impaired expression of E-cadherin *in vivo* (***: *p* < 0.001).

Several factors can be proposed as causative of the downregulation of E-cadherin in *BRAF*^mut^ CRC. One promising candidate is epigenetic DNA methylation because *BRAF* mutation in CRC is categorized within CMS1, which shows various DNA hypermethylation [5]. Kanazawa et al. evaluated the loss of heterozygosity in CRC and reported that promoter hypermethylation is a key epigenetic event in the loss of E-cadherin expression [24]. Currently, *SEPT9* and *NDRG4*/*BMP3* are approved as methylation-based diagnostic biomarkers for CRC [25]. A previous cohort study suggested that *CDH1* is a candidate methylation-based diagnostic biomarker for CRC [26]. We analyzed the PanCancer Atlas cohort data in TCGA (*N* = 312) and found that multiple CpG sites in the *CDH1* gene [27] showed higher levels of DNA methylation in *BRAF*^mut^ CRC than in *BRAF*^WT^ CRC (***: *p* < 0.001, **Figure 5c**). These findings indicate that epigenetic DNA methylation is one of the key factors causing impaired E-cadherin expression in *BRAF*^mut^ CRC *in vivo*.

### 2.6. Inhibition of DNA methylation partially reverse dynamic phenotype of *BRAF*^mut^ organoid

To verify our hypothesis that epigenetic DNA methylation is a factor causing impaired E-cadherin expression in *BRAF*^mut^ CRC, we treated CRC organoids with a methylation inhibitor, 5-aza-2’-deoxycytidine (5-azadc, Method S2, Supporting Information). As 5-azadc is incorporated into nucleic acids and prevents methylation at CpG sites via covalent bonding, it can be used to screen epigenetically masked genes in CRC [28]. The qRT-PCR data of *BRAF*^mut^ organoid (C7T) treated with DMSO (control) or 1 and 3 µM 5-azadc showed that the expression of CDH1 increased with increasing 5-azadc concentration (**Figure 6a**). The viscoelastic relaxation of Γ(*t*) of C7T treated with 3 µM 5-azadc and DMSO (control) implied that the viscoelastic response of C7T (*BRAF*^mut^) was accelerated by inhibiting DNA methylation (**Figure 6b**). The mean characteristic relaxation times in the absence and presence of 5-azadc are 〈*τ*_5−azadc_〉 ≈ 0.2 h and 〈*τ*_control_〉 ≈ 0.8 h, respectively (**Figure 6c**), suggesting that methylation inhibition had a significant effect on the viscoelasticity. In fact, the statistically significant increase in effective elasticity *k* (**Figure 6d**) and the significant decrease in effective viscosity *η* (**Figure 6e**) provided supporting evidence that DNA methylation affects the mechanical phenotypes of *BRAF*^mut^.

**Figure 6.**
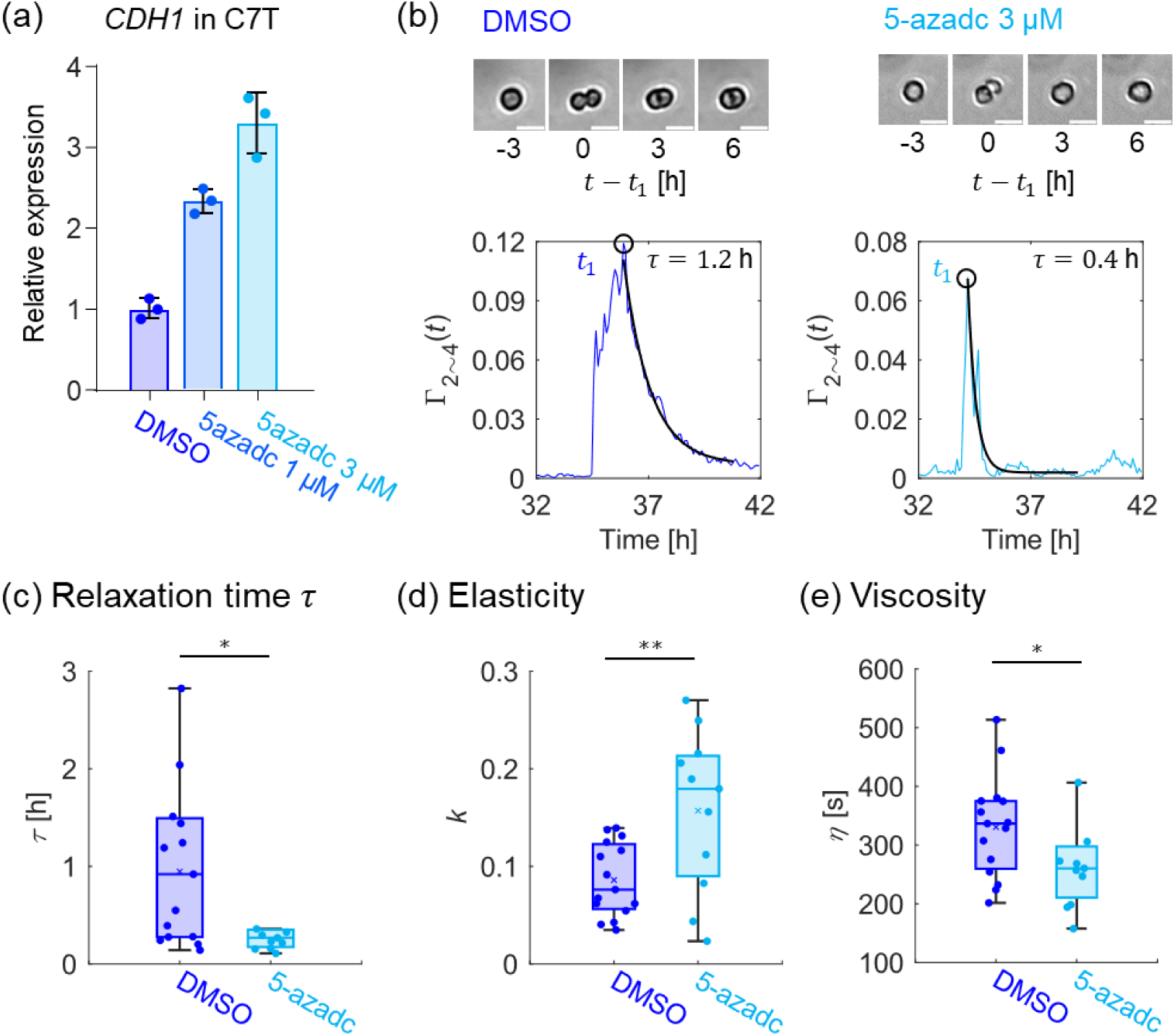
Methylation inhibitor partially inverts *CDH1* expression and dynamic phenotypes. (a) Expression level of *CDH1* in C7T treated with methylation inhibitor 5-azadc increases in a concentration-dependent manner. (b) C7T treated with 3 µM 5-azadc (light blue) exhibited much faster recovery than the control (blue), which was confirmed by the results of relaxation time *τ* (c), effective elasticity *k*(d) and viscosity *η* (e). DMSO, *N* ≥ 14; 5-azadc, *N* ≥ 9. *: *p* < 0.05, **: *p* < 0.01. Scale bars: 20 µm.

## 3. Discussion

Spatiotemporal analysis of patient-derived CRC organoids suggested that the mechanical properties relating to the adhesion between cells changed already at an early phase of organoid development, well ahead of the formation of hollow cysts (*t* ≤ 80 h). In contrast to widely taken approaches using image-based high-throughput screening of mature CRC organoids, our approach enables detection of the changes in viscoelasticity of patient-derived CRC organoids at an earlier timepoint in a noninvasive manner. Fourier mode analysis of 2D-projected images revealed that *BRAF*^mut^ organoids undergo more significant deformation than their wild-type equivalents. To date, morphological changes of 3D multicellular systems over time have been studied in the context of morphogenesis within various frameworks, such as the widely employed 3D vertex model [29] and analytical models that consider the balances between junctional tension and hydrostatic pressure [30].

In this study, organoids were treated as one continuum object, because we focused on the early phase (*t* = 0 ∼ 80 h), when they consisted of 8 ∼ 16 cells adopting a quasi-spherical geometry with no hollow cysts. Therefore, we employed a simple mathematical model considering cellular deformation and rearrangement as the *L*^2^-gradient flow of free energy in 2D space. Our model successfully reproduced the experimentally observed shape relaxation, highlighting that the intercellular tension and hence cell–cell contact cause the viscoelasticity of multicellular systems. While the application of physical concepts to complex, biological phenomena has considerably deepened our understanding in a variety of contexts [31], the extent to which this has been connected to the processes occurring at the molecular level remains limited. Thus, to obtain more insight into this issue at the molecular level, we compared the expression levels of intercellular E-cadherin, the main constituent of adherens junctions, and cortical F-actin, which generates the contractile forces together with myosin motor. Our data demonstrated the decrease in the expression of E-cadherin and cortical actin at the protein level, and the decrease in expression of the *CDH1* gene was further confirmed at the mRNA level by qRT-PCR.

Although E-cadherin was proposed as an additional biomarker for CRC to supplement the currently used biomarkers such as carcinoembryonic antigen (CEA) [20], apparently conflicting data suggesting different mechanisms behind the involvement of E-cadherin in CRC were reported [18]. Notably, many studies have shown that the redistribution of E-cadherin from membrane to cytoplasm plays more critical roles in CRC rather than its total expression level [32]. Khoursheed et al. reported that the level of membranous E-cadherin was generally significantly lower in tumor tissues than in their non-cancerous equivalents (*p* = 0.004), but this decline was not correlated with Duke’s staging, tumor grade, sex, or size and site of the tumor [33]. Meanwhile, Kim et al. reported that the loss of *CDH1* expression is associated with an infiltrative tumor growth pattern and lymph node metastasis but not significantly associated with prognosis [34]. Several studies have even suggested that E-cadherin can act not only as a tumor suppressor but also as a tumor promoter. For example, the increase in E-cadherin expression was observed in chemoresistant CRC cells [35], and E-cadherin-mediated intercellular junctions and cell polarity acted against Fas receptor-induced apoptosis [36]. It is known that E-cadherin expression can be altered via both genetic and epigenetic modifications, one of which is the epigenetic methylation of DNA. In CRC, several studies showed that frequent CpG island methylator phenotype (CIMP-H) is closely associated with both *BRAF* mutation and *MLH1* methylation, leading to high microsatellite instability (MSI-H) status [37]. Zheng et al. analyzed the relationship among mutation, expression and methylation levels in a PanCancer Atlas cohort data in TCGA in the context of CRC [38]. Among the frequently mutated genes, mutations in *KMT2C*, *CDH1*, *APC*, *PIK3CA*, *PTCH1* and *TP53* significantly affected their gene expression levels, while *BRAF*, *BRCA2*, *CDH1*, *NF1*, *PTCH1*, *SMO* and *TSC1* gene mutations affected their methylation levels. The partial recovery of dynamic phenotypes by 5-azadc treatment (**Figure 6**) indicated that epigenetic DNA methylation in *BRAF*^mut^ organoids is indeed a key regulator of impaired E-cadherin expression, resulting in the characteristic mechanical phenotype.

One of the critical signaling pathways associated with E-cadherin is canonical WNT3/β-catenin signaling pathway. In the context of CRC, nuclear translocation of β-catenin, a key protein in WNT signaling pathway, is a hallmark of tumor development and progression [39]. For example, mutations of *APC* that is critical for the formation of β-catenin destruction complex, often lead to the nuclear translocation of β-catenin. The immunohistochemical images of β-catenin in *BRAF*^WT^ and *BRAF*^mut^ CRC tissues (Figure S5, Supporting Information) suggest weaker nuclear β-catenin signals in *BRAF*^mut^ than in *BRAF*^WT^. This seems to agree well with the previous immunohistochemical study reporting lower nuclear β-catenin expression in the sessile serrated lesion and carcinoma with *BRAF* mutation [40]. The TCGA data showed significant suppressions of *WNT3A* (***: *p* < 0.001) and *CTNNB1* (**: *p* < 0.01) in *BRAF*^mut^ CRC (Figure S6, Supporting Information). It should be noted that the expression level of *WNT3A* was very low, close to the detection limit. Notably, both the upstream receptor of canonical WNT signaling, *FZD7,* as well as the downstream genes, *AXIN2* and *ASCL2*, were significantly downregulated (all showed ***: *p* < 0.001, Figure S6, Supporting Information). These data suggest that whole canonical WNT signaling is attenuated in *BRAF*^mut^ CRC compared to *BRAF*^WT^ CRC studied in this research. Mutations of WNT-related genes are common in serrated pathway, which leads to lower WNT activity [41]. The TCGA data analysis showed that 80% of *MLH1*-methylated CIMP-H CRC is characterized by *BRAF*^mut^ and MSI-H, as well as by downregulation of canonical WNT [42]. In turn, EGFR-mediated MAPK/MEK pathway is activated, which has been shown to cause the resistance of *BRAF*^mut^ CRC to RAF inhibitor [43]. This seems to be in line with our pathway analysis data that suggest the upregulation of PCP pathway (**Figure 4e**), which could explain mesenchymal, invasive characteristics of *BRAF*^mut^ CRC. These data suggest that hypermethylation of *CDH1* and downregulation of canonical WNT upregulate PCP pathway in *BRAF* mutated CRC, resulting in a significant modulation of viscoelasticity of early-stage organoids.

Remarkably, the mechanical properties, immunohistochemical signal intensity in organoids, distribution of E-cadherin in CRC tissues exhibited no significant difference between *KRAS*^mut^ CRC and the corresponding wild type (Figure S7, Supporting Information). *KRAS* mutation is common in the subgroup that is characterized by a low number of DNA methylation markers, called CIMP-L or CIMP2, leading to microsatellite stability (MSS) [37b, 44]. In contrast to *BRAF*^mut^ CRC (**Figure 5b**), TCGA data analysis showed that the difference in expression of *CDH1* in *KRAS*^mut^ and *KRAS*^WT^ was less significant compared to *BRAF* (*: *p* < 0.05, Figure S7f). In fact, CpG sites in the *CDH1* gene showed no significant difference between *KRAS*^mut^ and *KRAS*^WT^, suggesting that the methylation of *CDH1* plays no critical roles.

## 4. Conclusion

The present study focused on how driver gene mutation pattern is associated with the force-driven active deformation of patient-derived CRC organoids at an early phase (*t* ≤ 80 h). With aid of Fourier mode analysis, we quantitatively demonstrated that *BRAF*^mut^ organoids possess markedly lower elasticity and higher viscosity compared to *BRAF*^WT^ organoids. The mathematical model numerically indicated that the intercellular tension is the factor causing the change in viscoelastic shape recovery. On the molecular level, immunohistochemical staining of *BRAF*^mut^ and *BRAF*^WT^ organoids revealed a substantial difference in the expression of E-cadherin and F-actin, which are responsible for cell-cell adhesion and traction force generation, respectively. The downregulation of E-cadherin gene (*CDH1*) in *BRAF*^mut^ organoid was further confirmed on the mRNA level using qRT-PCR. Of note, the differential expression of E-cadherin between *BRAF*^mut^ and *BRAF*^WT^ in *in vivo* tissues was also confirmed by immunohistological staining of the original CRC tissues and TCGA cohort data, indicating that epigenetic DNA methylation is a key factor downregulating *CDH1*. This was verified by the concentration-dependent restoration of *CDH1* expression under the treatment with DNA methylation inhibitor 5-azadc. Remarkably, the treatment with 5-azadc also reverted the dynamic phenotype of *BRAF*^mut^ organoids to that of *BRAF*^WT^. These findings were further supported by cohort and pathway analyses, indicating the downregulation of the canonical WNT and upregulation of PCP, which could explain invasive and metastatic phenotypes of *BRAF*^mut^ CRC. As these physical indices, such as shape relaxation time and viscoelasticity, do not have any overlap with the information at the molecular level, the complementary combination of physical and biochemical approaches could help us unravel the causality connecting critical genetic and epigenetic alterations, pharmacological interventions, and prognostic phenotypes.

## 5. Methods

### 5.1. Patient-derived human CRC organoids

All CRC samples used to establish organoids were obtained from Kyoto University Hospital with written informed consent. The study protocol was approved by the Ethics Committee of Kyoto University Hospital (G1288). None of the patients had undergone chemotherapy or preoperative radiation. Surgically resected CRC tissues were washed with PBS and minced using scissors, and digested in 0.2% collagenase type I (Thermo Fisher Scientific) at 37°C for 60 min. The dissociated cells were embedded in 25 µL of Matrigel (Corning) and cultured in advanced DMEM (Gibco) supplemented with 1 % penicillin–streptomycin (Sigma), Glutamax (Gibco), 5 % inactivated FBS (Gibco), 10 μM Y-27632 (TOCRIS), 1 μM SB431542 (TOCRIS), 50 ng/mL EGF (PEPRO) and 100 ng/mL b-FGF (PEPRO). Organoids were passaged by dissociation with 1 mL of TrypLE Express (Gibco) for 10 min at 37 °C. Following the addition of 4 mL of medium and centrifugation at 300 × g, cells were reseeded in Matrigel and covered with organoid medium. The medium was refreshed every other day, and organoids were passaged every week.

### 5.2. Immunohistochemical imaging

Whole-mount immunostaining of organoids was conducted in accordance with a previous report [45]. In short, Matrigel was dissociated in a cell recovery solution (Corning) for 60 min at 4 °C. At *t* = 72 ∼ 80 h, organoids were fixed in 4 % paraformaldehyde (PFA) at 4 °C for 45 min and then permeabilized with 0.1 % Tween-20 for 10 min. After blocking with wash buffer (0.2 % BSA, 0.01 % SDS, and 0.1 % Triton X-100), the organoids were incubated at 4 °C overnight with primary antibodies. After washing the primary antibody, the organoids were incubated overnight with the secondary antibodies at 4 °C. Immunofluorescence images were acquired using an A1R confocal laser scanning microscope (Nikon, Japan). Immunostaining of tissues was conducted after the tissues had been fixed in masked formalin 2A (Japantanner), embedded in paraffin and cut into 5-μm-thick sections. The sectioned samples were deparaffinized, rehydrated and treated for antigen retrieval. After incubation in protein block serum-free solutions (DAKO), tissues were incubated with the primary antibody overnight at 4 °C, followed by incubation with secondary antibody. Further details can be found in Method S3, Supporting Information.

### 5.3. Live imaging and image analysis

Time-lapse imaging of the active deformation of organoids growing from the single-cell stage was performed using an IX81 inverted microscope (Olympus, Japan) equipped with a humidified, temperature-controlled chamber containing an atmosphere with 5 % CO_2_. For each set of experimental conditions, 6–10 x-y positions and 3–5 z positions (per x-y point) per well were selected. Brightfield images were acquired every 10 min using a 10× objective lens (Olympus, Japan). The time-lapse images of organoids were segmented using Cellpose. A montage of 10 × 10 time-lapse images was subjected to the analysis. After segmentation, spatiotemporal dynamics were analyzed using a custom-made algorithm in MATLAB (MathWorks).

### 5.4. qRT-PCR and RNAseq

Total RNA was extracted using RNA Mini Kit (Qiagen, Hilden, Germany). Single-stranded complementary DNA (cDNA) was synthesized using ReverTra Ace qPCR RT Kit (TOYOBO, Tokyo, Japan). qRT-PCR was performed using SYBR Green Master Mix (Roche Applied Science) and LightCycler 480 (Roche Applied Science). The expression levels were standardized by comparison with those of GAPDH. RNAseq was performed by Macrogen, Inc. (Seoul, South Korea), on the NovaSeq6000 platform with 2 × 100 bp paired-end sequencing using the TruSeq RNA Sample Prep Kit v2. Adaptors and low-quality bases were trimmed from the reads using Trimmomatic 2 (version 0.39) with the default parameters. Reads were mapped to the reference genome using STAR software (version 2.7.3a) and counted using RSEM (version 1.3.1).

### 5.5. Statistical Analysis

Statistical analysis was performed using in MATLAB (MathWorks). All box plots presented in this study present the median value as a solid line and the average value as a cross mark. The boxes correspond to the interquartile ranges (IQR), and the whiskers extend to the most extreme values within 1.5 × *IQR* from first (*Q*_1_) and third (*Q*_3_) quartiles, where *IQR* = *Q*_3_ − *Q*_1_. Comparisons between two groups were performed using Student’s *t* test. The *p* values < 0.05 were considered as significant difference. Outliers were identified using the IQR method [46], where any value beyond the whiskers was considered an outlier.

## Acknowledgements

S.N., R.S. and M.T. thank Satoko Hinatsu for experimental assistance and Kentaro Hayashi for useful comments on the immunofluorescence image analysis. M.T. thanks Thomas Holstein and Martin Bastmeyer for critical comments, and S.N. thanks Karel Svadlenka and Tetsuya Hiraiwa for insightful suggestions on the mathematical modelling. This study was supported by JSPS KAKENHI (JP20H00661 to M.T. and H.S., JP24H00796 to M.T., and JP22H03939 to R.S.) and the German Science Foundation (Germany’s Excellence Strategy—2082/1—390761711 and CRC1324 A05 to M.T.). M.T. also thanks the Nakatani Foundation for its support, and S.N. thanks DAAD for the PhD fellowship. The authors thank German-Japanese HeKKSaGOn Alliance for support and Scribendi (www.scribendi.com) for editing a draft of this manuscript.

## Data Availability Statement

The authors declare that the main data supporting the results in this study are available within the paper and its Supplementary Information. The raw and analyzed datasets generated during the study are available for research purposes from the corresponding authors (M.T. and H.S.) on reasonable request.

## Author Contributions

Conceptualization and supervision: M.T., H.S. and A.F.; live imaging, spatiotemporal analysis, mathematical modelling, and immunofluorescence imaging: S.N. and R.S.; RNAseq, qRT-PCR, TCGA analysis: G.Y.; writing–original draft: M.T., S.N. and G.Y.; reviewing and editing: R.S., A.F., H.S. and M.T. All authors proofread, commented on and approved the final version of this manuscript.

## Supporting Information

Supporting Information is available from the author.

## Table of Contents

Analyses of dynamic tissue deformation and mathematical modeling reveal the epigenetic downregulation of intercellular adhesion in colorectal cancer organoids with *BRAF* mutation. Recovery of cadherin expression by pharmacological treatment can be detected as modulation of viscoelasticity, suggesting that the combination of dynamic phenotyping, modeling, and molecular level analyses can unravel the mechanistic causality connecting gene mutations, therapeutic interventions, and clinical outcomes.

