## Supplementary material for "Spatiotemporal Patterns of Active Deformation Reveal Downregulation of Cell-Cell Adhesion in Patient-Derived Colorectal Cancer Organoids with *BRAF* Mutations": Supplental Data

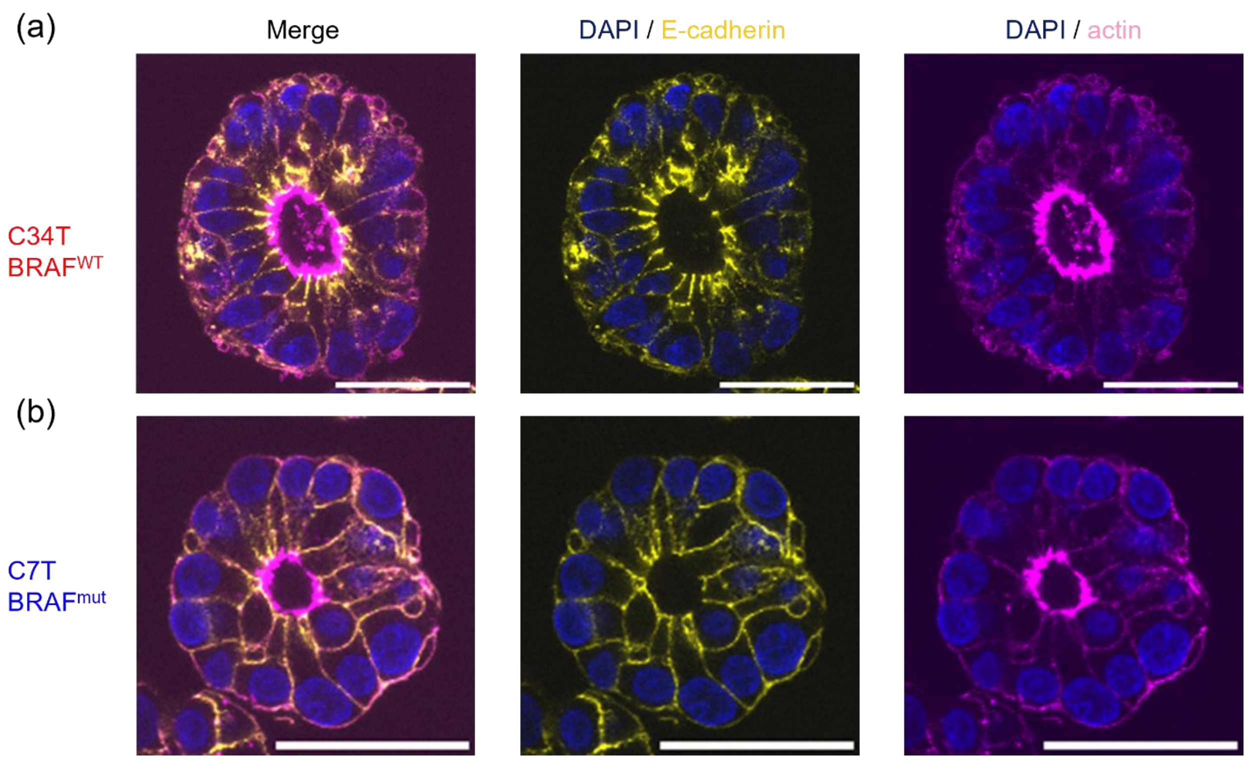


**Figure S1. Cyst formation of patient-derived colorectal cancer organoids on day 5**

Immunohistochemical images of E-cadherin (yellow) and F-actin (magenta) for $BRAF^{WT}$ (C34T, a) and $BRAF^{mut}$ (C7T, b). The organoids were fixed on Day 5. Each image is a slice corresponding to the plane of equator. The hollow cysts were formed in both $BRAF^{WT}$ and $BRAF^{mut}$. Note that $BRAF^{WT}$ organoid formed a cyst surrounded by a cell monolayer with the apico-basal polarity, while $BRAF^{mut}$ organoid did not form a monolayer. Scale bars: 40 μm.

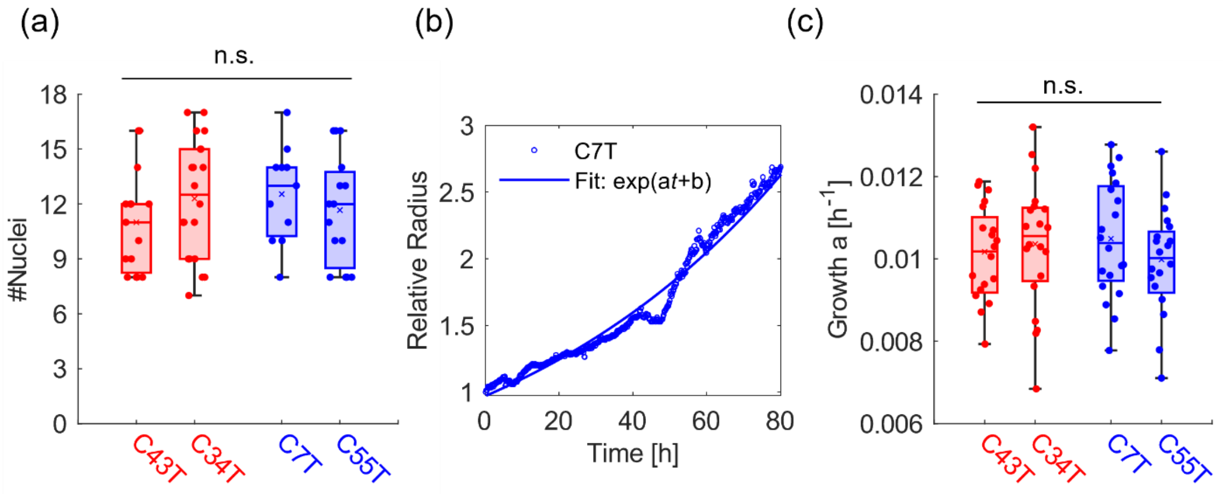


**Figure S2. Proliferation rate of organoids**

(a) The number of nuclei quantified from immunohistochemical images acquired at $t \approx 80$ h revealed no significant difference between $BRAF^{\mathrm{WT}}$ and $BRAF^{\mathrm{mut}}$ organoids. (b) Time-course of increase in radius of a C7T organoid (normalized by the initial size) was well-fitted by an exponential function, $\exp(at + b)$. (c) Statistical comparison of the growth rate constant $a$, obtained from the exponential fitting, showed no significant difference between $BRAF^{\mathrm{WT}}$ and $BRAF^{\mathrm{mut}}$ organoids.

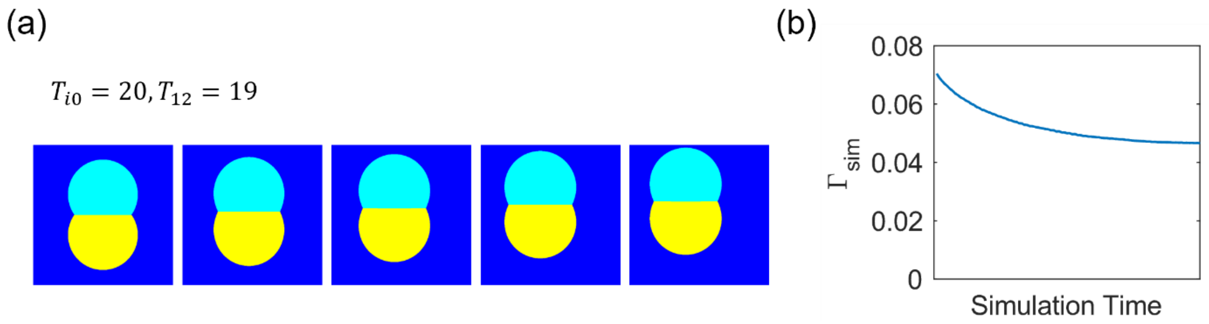


**Figure S3. Simulation with a higher intercellular tension $T_{12} = 19$**

(a) Snapshot images of simulation with intercellular tension $T_{12} = 19$, which is very close to the tension between cells and their surroundings ($T_{i0} = 20$). (b) Time-course of calculated deformation $\Gamma_{\text{sim}}$ corresponding to (a), indicating that cells hardly recover the circular shape.

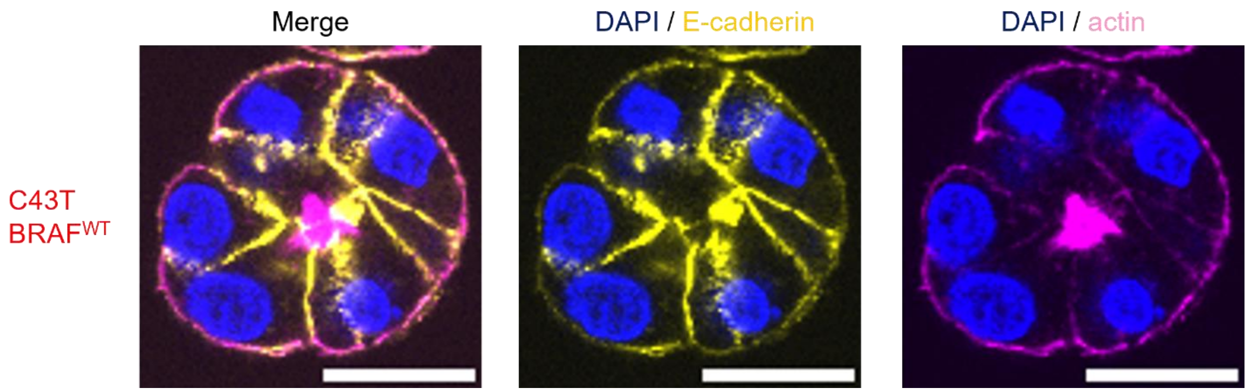


**Figure S4. Condensation of actin near the future cyst**

The immunohistochemistry images of E-cadherin (yellow) and F-actin (magenta) show the condensation of actin near the centre, corresponding to the position of future cyst. Scale bars: 20 μm.

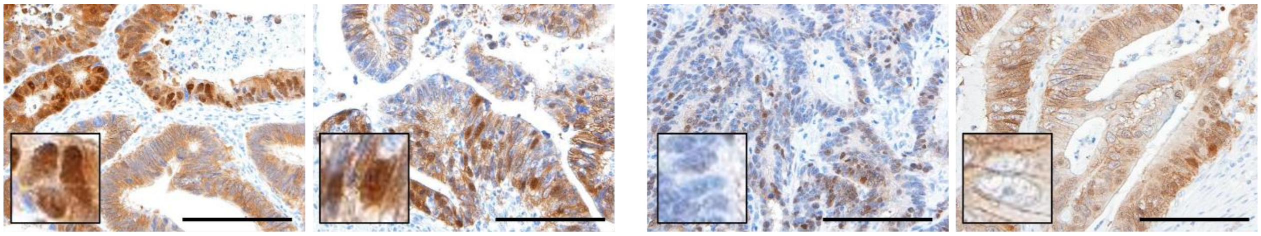


**Figure S5. Immunohistochemical images of β-catenin**

Immunohistochemical images of β-catenin of original CRC tissues for *BRAF*$^{WT}$ (C43T, C34T) and *BRAF*$^{mut}$ (C7T, C55T). Higher-magnification images suggest that tissues with *BRAF*$^{mut}$ have poorer nuclear signals compared to those with *BRAF*$^{WT}$. Scale bars: 100 μm.

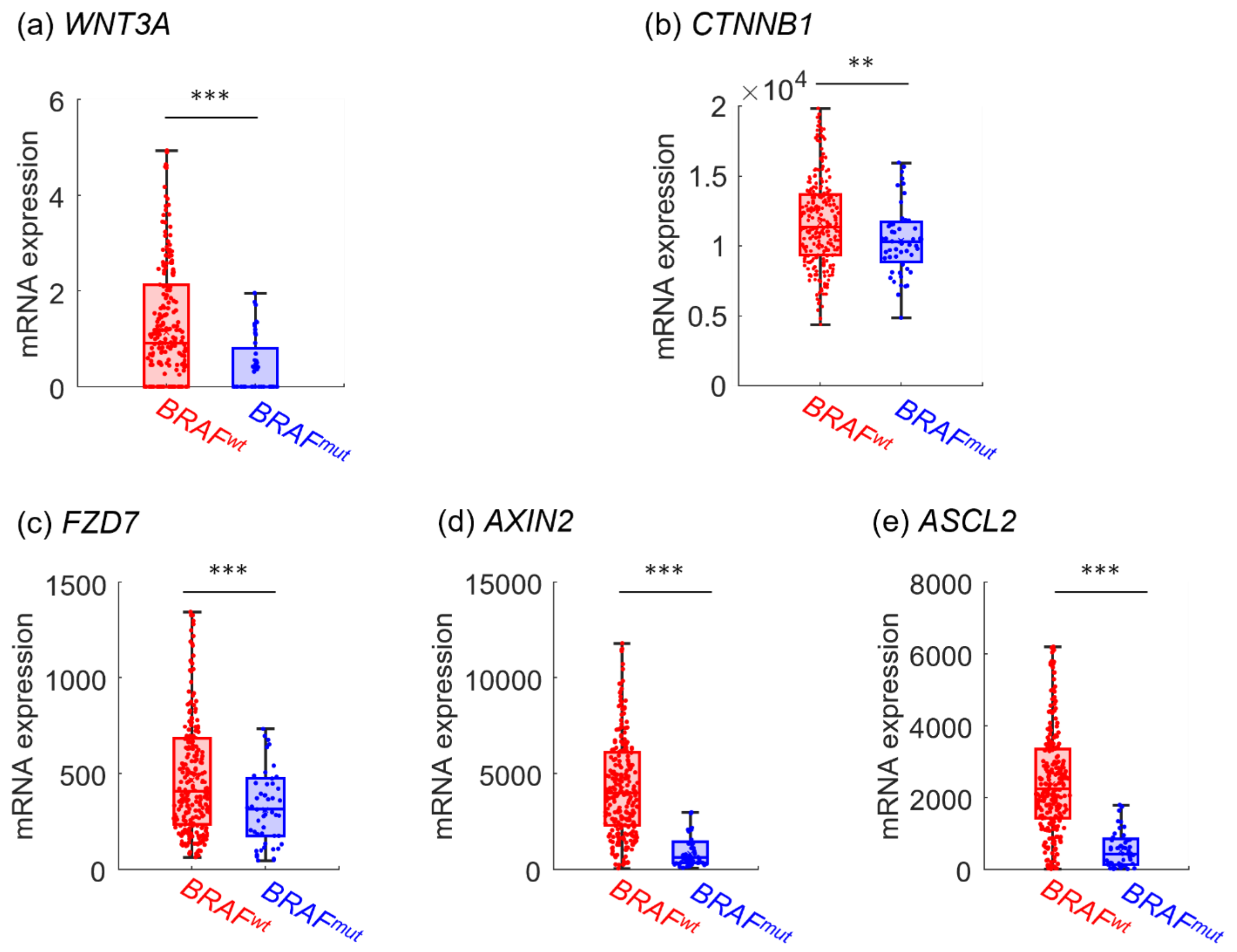


**Figure S6. Modulation of Wnt and Wnt-associated genes**

PanCancer Atlas cohort from TCGA shows significant decrease in *WNT3A* and *CTNNB1* expression of *BRAF*$^{\mathrm{mut}}$ CRC tissues compared to *BRAF*$^{\mathrm{WT}}$. Notably, *BRAF*$^{\mathrm{mut}}$ significantly downregulate the upstream gene, *FZD7* and downstream genes, *AXIN2* and *ASCL2*, of canonical Wnt pathway ($** : p < 0.01,\ *** : p < 0.001$).

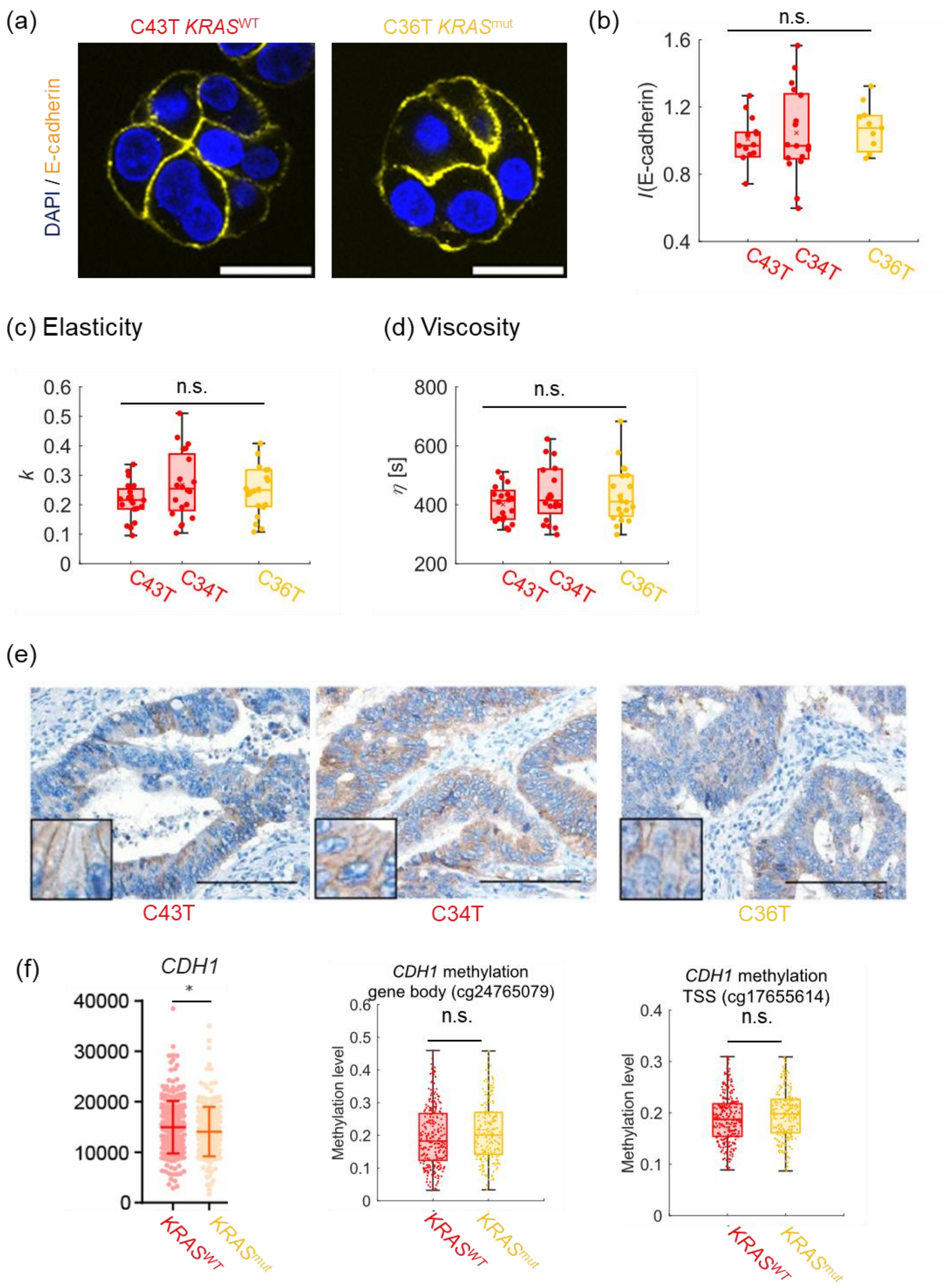


**Figure S7. Mechanical properties and expression of *CDH1* and E-cadherin in *KRAS*$^{\mathrm{mut}}$ and *KRAS*$^{\mathrm{WT}}$ organoids**

(a) Immunohistochemical images of E-cadherin (yellow) in *KRAS*$^{\mathrm{WT}}$ (left) and *KRAS*$^{\mathrm{mut}}$ (right) organoids fixed on Day3. Scale bars: 20 μm. (b) Statistical comparison of integrated E-cadherin signals shows no significant difference between *KRAS*$^{\mathrm{WT}}$ (C43T, C34T, red) and *KRAS*$^{\mathrm{mut}}$ (C36T, yellow) organoids. (c, d) The calculated elasticity and viscosity exhibit no significant differences between *KRAS*$^{WT}$ (C43T, C34T, red) and *KRAS*$^{mut}$ (C36T, yellow). (e) Immunohistological images of E-cadherin in *KRAS*$^{WT}$ (C43T, C34T) and *KRAS*$^{mut}$ (C36T) tissues. Scale bars: 100 μm. (f) Expression profile and the methlyation levels of *CDH1* obtained from TCGA datasets showed much less difference between *KRAS*$^{WT}$ and *KRAS*$^{mut}$.

**Method S1. Numerical simulation for relaxation of organoid**

For Numerical realization, level set-based methods were employed. Mainly, the heat equations for the boundaries between regions were solved, and level sets of their solutions were extracted. To preserve the area of each region ($C_i$), auction dynamics were combined with level set-based methods.

The intercellular line tension $T_{12}$ was set 1, 12, and 19, and the line tension between cells and their surroundings $T_{0i}$ was set 20. The simulation domain [0,1] × [0,1] was uniformly discretized into 512 × 512 points with periodic boundary conditions. The time step was set 0.001, and 100 time steps were implemented. The initial condition on a simulation was shown in Figure 3a.

**Method S2. Pharmacological inhibition of DNA methylation.**

The CRC organoid with *BRAF* mutation (C7T) was treated with DMSO (control) or with the demethylating agent 5-azadc (Selleck, Cat# S1200) at concentrations of 1 μM and 3 μM for 96 h, and subjected to the qRT-PCR analysis or time-lapse imaging.

**Method S3. Whole-mount immunohistochemical staining of organoids**

Whole-mount immunohistochemical staining was performed on organoids using an anti-E-cadherin primary antibody (ab1416, Abcam), diluted at 1:400. A secondary antibody, Goat anti-Mouse IgG (H+L) Highly Cross-Absorbed Secondary Antibody, Alexa Fluor$^{TM}$ Plus 488 (A32731, Thermo Fisher) was applied at a final dilution of 1:1000. Nuclei and F-actin were stained with DAPI (1:5000) and phalloidin (1:1000), respectively. For the quantitative comparison of integrated E-cadherin and F-actin signals across multiple experiments, organoids derived from the C34T ($BRAF^{WT}$) were used as an internal control.

**Movie S1. Time-lapse movie for *BRAF*$^{\mathrm{WT}}$ organoid from single-cell stage up to $t \approx 80$ h (scale bar: 20 μm)**

**Movie S2. Time-lapse movie for *BRAF*$^{\mathrm{mut}}$ organoid from single-cell stage up to $t \approx 80$ h (scale bar: 20 μm)**

**Movie S3. Numerical simulation with intercellular line tension $T_{12} = 1$**

**Movie S4. Numerical simulation with intercellular line tension $T_{12} = 12$**

**Movie S5. Numerical simulation with intercellular line tension $T_{12} = 19$**
